# Automated Raman micro-spectroscopy of epithelial cells for the high-throughput classification

**DOI:** 10.1101/2021.04.23.441084

**Authors:** Kevin O’Dwyer, Katarina Domijan, Adam Dignam, Marion Butler, Bryan M. Hennelly

## Abstract

Raman micro-spectroscopy is a powerful technique for the identification and classification of cancer cells and tissues. In recent years, the application of Raman spectroscopy to detect bladder, cervical, and oral cytological samples has been reported to have an accuracy that is greater than standard pathology. However, despite being entirely non-invasive and relatively inexpensive, the slow recording time, and lack of reproducibility, have prevented the clinical adoption of the technology. Here we present an automated Raman cytology system that can facilitate high-throughput screening and improve reproducibility. The proposed system is designed to be integrated directly into the standard pathology clinic, taking into account their methodologies and consumables. The system employs image processing algorithms and integrated hardware/software architectures in order to achieve automation and is tested using the ThinPrep standard, including the use of glass slides, and a number of bladder cancer cell lines. The entire automation process is implemented using the open source Micro-Manager platform, and is made freely available. We believe this code can be readily integrated into existing commercial Raman micro-spectrometers.

## 1 Introduction

Raman micro-spectroscopy (RMS) is a powerful technique for the identification, classification, and diagnosis of cancer cells and tissues. [1–9] The prefix “micro” indicates integration of a Raman spectroscopy (RS) system into a microscope, such that spectra are obtained from sub-micron areas in a sample. RS is based on an inelastic light-matter scattering, whereby the scattered photons have lost energy relative to the incident photons equivalent to the energy of certain Raman-active molecular bonds present within the sample. RS can identify biomolecular changes within cells as they progress from a healthy to a cancerous state. [1–3] The magnitude of the frequency shift is dependent on the molecular structure of the sample. For a complex mixture of different chemicals, such as in a cell, a highly unique Raman spectrum can be observed that is sometimes described as a ‘fingerprint’. Multivariate statistical analysis is applied to Raman spectra for classification, whereby statistical pattern recognition algorithms, such as Principal Components Analysis (PCA) combined with Linear Discriminant Analysis (LDA), are applied to identify subtle changes across datasets that can be used to accurately differentiate between different pathological groups. [1–8]

Malignancies of epithelial tissue, or carcinoma, account for 80-90% of all cancer cases [10], and clinical cytology, whereby oral, cervical, or bladder epithelial cells are visually inspected, is routinely applied in pathology labs for diagnostics. Distortion of the cell nucleus is common for carcinoma, and pathologists are trained to identify pertinent morphological features. The applications of RMS to clinical cytology with specific targeting of the cell nucleus for classification has shown promising results with many reporting the classification of bladder, [3–6] cervical, [7,11–13] and oral [8,14] cytological samples with an accuracy that is significantly greater than standard pathology. However, issues around reproducibility and speed have thus far hindered the clinical adoption of Raman cytology into the clinic. In this paper, we focus on the development of an automated RMS system for the classification of epithelial cells, based on the biochemical composition of the nucleus, with the ultimate goal of high-throughput noninvasive screening.

This paper builds on recent work in the development of robust Raman cytology platforms that are fully automated and high-speed. Schie et. al. have published several papers that describe a high-throughput automated Raman cytology platform with several applications. [15–21] The basic platform [15,16] has several important features including a defocused laser spot to excite a large cell area with high power, thereby facilitating shorter exposure times, and a Labview-based automaton process, which identifies cell position using image processing and controls a translation stage to align the cells with the laser spot. The system employs a 400 mW laser with a wavelength of 785 nm focused to an area of 10 *μm*^2^, which enables acquisition times <1 s for various cell types. The image processing component is central to the overall system. This is based on several steps, which in essence, identify the cell position as the centre of the bulk cell area, appearing dark with respect to the bright-field background. Combined with multivariate statistical classification, the system has been demonstrated to automatically classify hundreds of thousands of different cells based on their Raman spectra in relatively short times. In its first application, the system was successfully applied to various white-blood cells as well as pancreatic cells [15,16] with a maximum rate of 5000 cells/hour. Drug testing is a key application area for automated Raman cytology systems. In this capacity, the same system has been applied by the authors to (i) study the effect of dithiothreitol on the diatom Phaedactylum tricornutum; [17] (ii) to analyse multi-well plates [18] for the purpose of developing a Raman-based cell viability assay; specifically, on the effect of doxorubicin concentration on monocytic THP-1 cells; and (iii) to investigate the effect of the targeted cancer drug panitumumab on colorectal cancer cell lines. [20]

Although, not a target application of the system proposed here, the study of pathogens, and in particular bacteria, has been another key application area of automated Raman cytology systems in recent years. The automated system described in the previous paragraph has been adapted to record spectra of single isolated neutrophils from human peripheral blood [19], which were stimulated via an in-vitro infection model with heat-inactivated bacterial and fungi pathogens; the system captured 20000 neutrophil spectra, across various treatment groups, originating from three donors. Another system developed by Douet et. al. [22] has been designed to provided automated Raman spectrocopy of individual bacteria cells; once again, this automated system is based on image processing in order to automatically identify the bacterial cell position, followed by alignment of the cell with the excitation laser. In this case, the image processing component relies on the availability of an out-of-focus diffraction pattern facilitated by the use of a spatially coherent illumination source. The recorded image can be described as an in-line digital hologram, which can be subjected to numerical propagation [23] in order to obtain an in-focus image of the sample. The cell position can be identified based only on image contrast, whereby the bacterial cell appears dark in a bright background.

In this paper, we present an automated Raman cytology system with several contributions: (1) This system utilises a simpler image processing component than previous systems, which is based on a single step. It is shown that this approach can accurately identify epithelial cell nucleus position, which to the best of our knowledge, has not been a target area for previous automated systems; (2) The system can be applied to unlabelled ‘phase-only’ adherent cells, which produce low image contrast; (3) The system is demonstrated to work with the ThinPrep standard, [5,7,12,24] an established instrument and protocol used to prepare cytology samples in hospital settings such as the cervical ‘Pap’ smear; (4) The method is based around the open-source Micro-Manager platform [25], which is freely available. The associated code is supplied in an online repository [26] and is described in detail in the supplementary information. This approach can be implemented easily on any existing RMS system that uses a motorized translation stage and a computer controllable microscope lamp and excitation laser, which are commonplace in modern life-science microscopes and commercial RMS systems.

## 2 Automation

### 2.1 Principle of Automation: Identifying Cell Nucleus Position using the Nucleus ‘Microlens-Effect’

Central to the proposed automated Raman cytology system is an image capture and processing methodology that facilitates the rapid identification of cell nuclei, which can subsequently be targeted for RS. The cell nucleus, which contains almost all of the cell DNA, is the primary target for Raman-based classification of epithelial cell type; [1–9,11–14] in some studies the nucleus is targeted at several different points and the resulting spectra are averaged, and in other studies, this ‘averaging’ is achieved optically by using a relatively large laser spot for excitation. However, this approach is complicated by the difficulty in clearly identifying cellular features such as the cell nucleus using brightfield microscopy. Adherent epithelial cells are commonly described as a weakly scattering phase-only objects that appear almost transparent when imaged using brightfield microscopy. Although several imaging modalities exist to improve the image contrast of such objects, such as phase-contrast, and differential interference contrast, these methods cannot reliably be used to identify the cell nucleus. Furthermore, these modalities require dedicated equipment such as a phase-contrast objective (which includes an annular filter) or polarising optics, which are preferably avoided when using RS. Fluorescence microscopy is the gold standard for identification of the cell nucleus, whereby a fluorescent stain such as 4’,6-diamidino-2-phenylindole (DAPI), which can penetrate the cell membrane and bind to adenine-thymine-rich regions in DNA. However, combining fluorescent microscopy and RMS is not straightforward, since both methods require dedicated excitation sources, and different optical collection systems and filters behind the microscope objective. The fluorescence spectrum can also overlap with the Raman spectrum although, it is possible to ensure that the two bands are mutually exclusive. [27] A further complication is that unlike RS, fluorescent labels are commonly incompatible with living cells. [28]

In this paper, we utilise the ‘microlens-effect’ of the cell nucleus in order to identify its position, whereby the phase-delay introduced by the cell nucleus has the effect of focusing partially-coherent illumination, similar to a effect of a micro-lens. This phenomenon has previously been investigated for the purpose of cell-counting, [29] which was the inspiration for this paper. In that study, human neuroblastoma cells were imaged in culture using a low magnification objective in order to capture a large number of cells in the field of view. The authors demonstrated that a green filter and a 130μm pinhole placed immediately above the culture flask generated bright-spots in a defocused plane ≈50μm from the object plane, that were approximately centred in the fluorescing regions (nuclei) of the cells. This was confirmed using the green fluorescent protein-tagged nuclear histone H2B protein. In the system proposed in this paper, partially coherent illumination is generated using only the microscope components that would be found in an existing RMS system. Such illumination can easily be obtained by closing the condenser aperture diaphragm such that the spatial coherence of the Kohler-illumination is maximised. We also make no attempt to apply a colour filter to the Tungsten Halogen lamp in order to enhance the bright-spot contrast. Another important difference in our work with respect to [29] is that the approach taken here makes use of a high magnification/numerical aperture microscope objective, which can resolve several bright spots in the ‘focal-plane’ of the cell nucleus corresponding to various sub-cellular features. Even with this simplified approach, we demonstrate that it is possible to identify the cell nucleus position with a sufficient degree of accuracy for subsequent targeting with RMS.

The applicability of this method of nuclear targeting is demonstrated for two forms of sample preparation: (1) live adherent epithelial cells in medium as shown in Fig.1, and (2) the same cells prepared using the ThinPrep standard as shown in Fig.2. ThinPrep is a clinical standard for the preparation of cyto-histological samples, particularly in the areas of cervical screening and urine cytology. The ThinPrep standard, including the use of associated fixatives and glass slides, has previously been shown to be compatible with Raman micro-spectroscopy. [5,7,12,24,30] HeLa cells were selected for this initial investigation due to their well known morphology and were prepared as described in Section 3.1.1 and imaged using an IX81 fluorescence microscope as described in Section 3.2. In Fig.1 (a) the live cells are shown in medium. The image was slightly defocused by displacing the sample approximately 1μm from the focal plane, and oblique illumination was used in order to improve visualisation of cell boundaries. In Fig.1 (b) the corresponding fluoresence image using DAPI is shown, which highlights the nucleus in each cell. In Fig.1 (c) the bright-spot image is shown. This was obtained by moving the sample up a distance of 50μm from the focal plane, which was found to provide the highest contrast image, although larger displacements could still be used to successfully resolve the bright spots. In this case, axial illumination is used to ensure the position of the bright-spot aligns with the nucleus of the cells when positioned in the image plane; In Fig.1 (d) the information from these three images is combined. Images (a) and (b) are superimposed together with the positions of the local maxima in image (c), which are shown as yellow targets. The red arrows highlight nuclei that have been just missed by the target. The green arrows highlight cells that appear to have been correctly targeted but did not fluoresce. The brown arrow highlight cases that are just at the edge of the nucleus but which would likely provide meaningful Raman spectra for a laser spot size > *1μm.* Finally, the orange arrows highlight cell nuclei that have been double targeted. Based on an analysis of several such images, we estimate successful (single) targeting of > 75% of HeLa cell nuclei.

**Fig 1.**
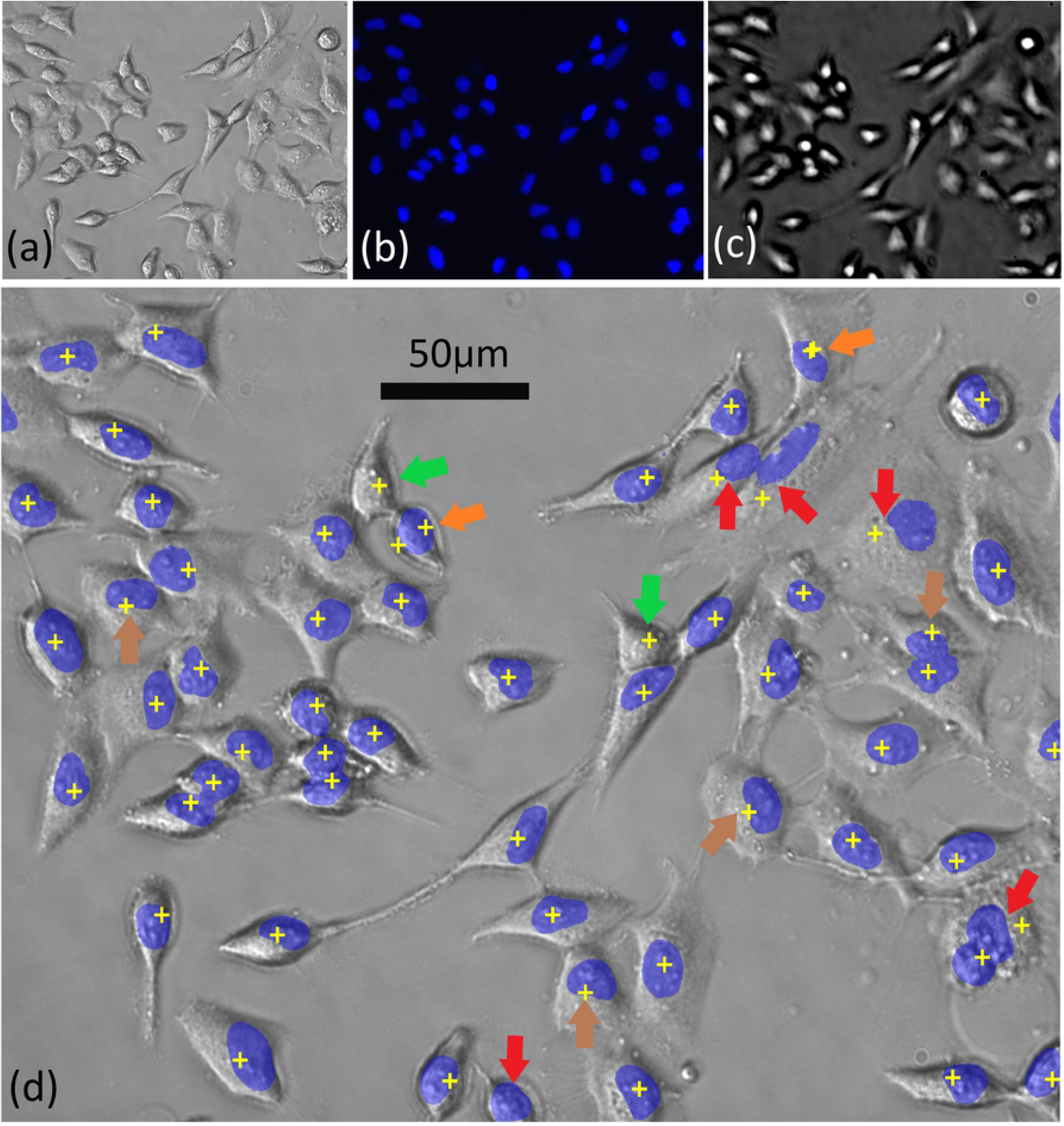
**Live HeLa cells in medium: (a) Brightfield image; (b) Fluorescence image using DAPI; (c) Bright-spot image: sample at** 50*μm* **from focal plane; (d) Combined images. Images (a) and (b) have been superimposed together with the positions of the local maxima in image (c) following image processing. The arrows are described in the text.**

**Fig 2.**
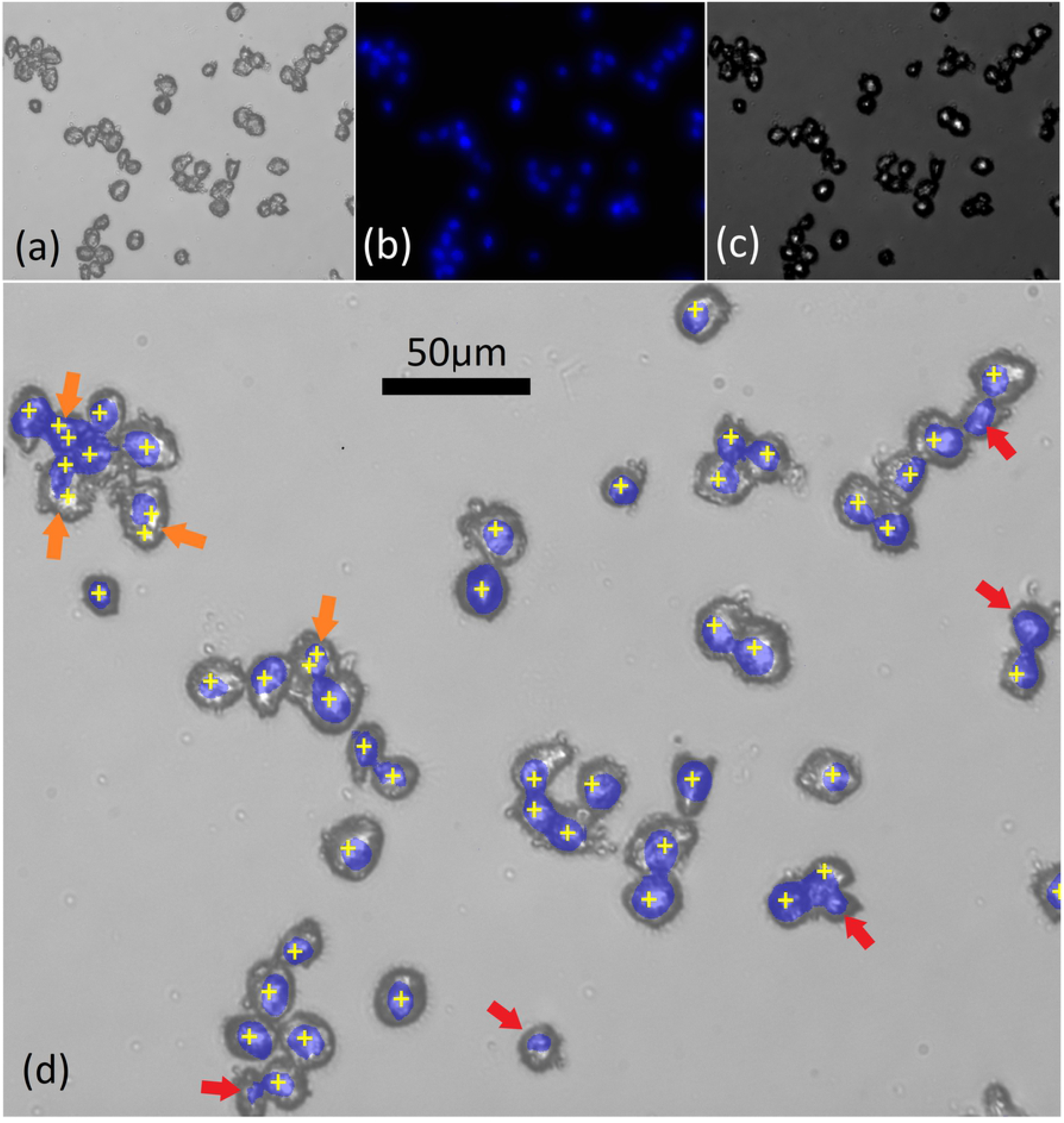
**HeLa cells on ThinPrep slides: (a) Brightfield image; (b) Fluorescence image using DAPI; (c) Bright-spot image: sample at** 14*μm* **from focal plane; (d) Combined images. Images (a) and (b) have been superimposed together with the positions of the local maxima in image (c) following image processing. The arrows are described in the text.**

A similar analysis was applied to samples prepared using the ThinPrep standard. A bright field image of these cells is shown in Fig. 2 (a). The cells have a thicker morphology and a greater depth of field than for the adherent case making it more difficult to record an in-focus image with a high NA objective. In Fig.2 (b) the corresponding fluorescence image using DAPI is shown, once again highlighting the nucleus in each cell. It is clear that the nuclei for the ThinPrep case are smaller in area than for the adherent case. In Fig.2 (c) the bright-spot image is shown. This was obtained by moving the sample up a distance of 14μm from the focal plane, which was found to provide optimal contrast. Interestingly, the ‘focal-length’ of the ThinPrep HeLa cells (i.e. the axial displacement providing the highest bright-spot image contrast) is significantly shorter than for the case of adherent cells, owing to their thicker rounder morphology. In Fig.2 (d) the information from these three images is combined. Images (a) and (b) are superimposed together with the positions of the local maxima in image (c) following Gaussian filtering, which are shown as yellow targets. The red arrows highlight nuclei that have been just missed by the target and the orange arrow highlight cases that are double targeted. Based on an analysis of several such images, we estimate successful (single) targeting of > 85% of HeLa cell nuclei prepared using the ThinPrep standard.

In this section, we have demonstrated that it is possible to identify cell nucleus position using a standard brightfield microscope with a closed condenser aperture, both for adherent live cell and also for cells prepared using the ThinPrep standard. In the next section, we outline a global automation routine for Raman cytology that makes use of this. Although in subsequent sections we focus our results on ThinPrep samples due to their clinical relevance, we have also found that adherent cells work equally well with this automated routine.

### 2.2 Global Automation Process

The nucleus ‘microlens-effect’ is used as a basis for targeting of cells for an automated RMS platform. The system uses a conventional Olympus-IX81 microscope, controlled with PC via the IX2-UCB control box, which allows for electronic control of the microscope objective focus position (Z-stage) and the white-light lamp. As illustrated in Fig.3, the open-source Micro-Manager software system [25] can be used to control the IX2-UCB, as well as several other opto-electronic components in the system, including an inexpensive CMOS digital camera inserted into the eyepiece of the microscope, the XY-translation stage, the laser used for Raman excitation, and the cooled CCD detector used to capture the Raman spectrum, all of which are described in more detail in Section 3.3. Micro-Manager allows for automatic control of these components with a modified Beanshell scripting interface, which also facilitates the use of the ImageJ library for image processing [31,32]. The entirety of the automation platform for collection of the cell spectra is written using this scripting interface and these scripts are freely provided in an online repository [26]. A key feature of this automation system is that the condenser aperture is closed to a minimum in order to maximise the contrast in the bright-spot image as described in the previous section.

**Fig 3.**
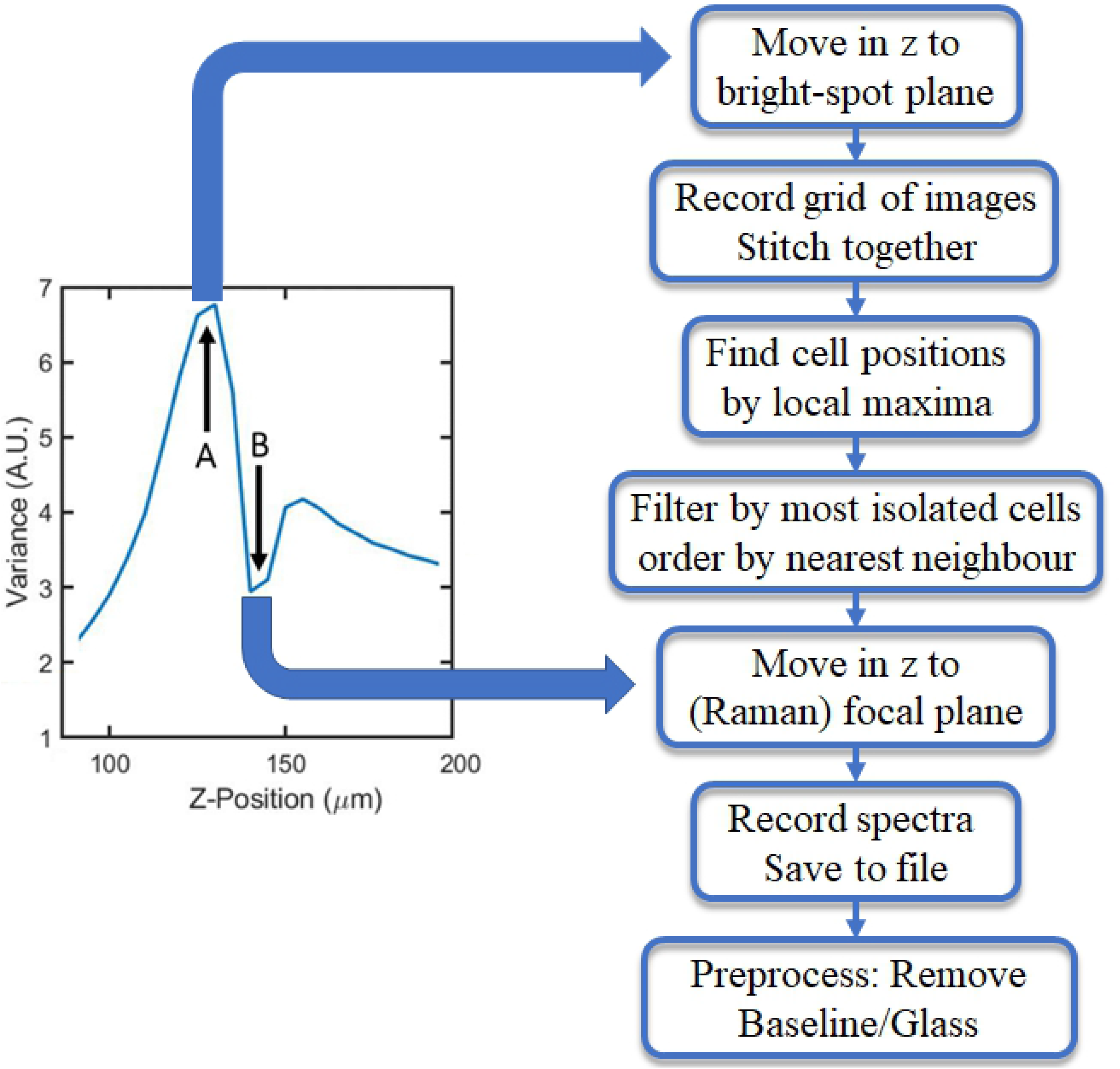
Schematic of opto-electronic system. The computer controls several elements in the system using an open-source integrated hardware control and image processing system, Micro-Manager. The various components, and symbols, are described in Section 3.3

In order to locate the cells, the Z-position of the bright-spot plane must first be determined. However, as described in the previous section, the position of this plane will vary significantly depending on cell morphology Therefore, in order to account for different cell morphologies, this plane is found using an auto-focusing routine. A series of spatially coherent light images are recorded across a range of focal positions in the same field-of-view (FOV). It was found that when the variance of a given FOV is maximised (defined as the square of the standard deviation over the mean for a given image’s pixel intensities), the sample is in the optimal bright-spot plane in terms of image contrast. The sample can also be brought into focus for brightfield imaging (and Raman spectroscopy) by finding the local minima of variance closest to the bright-spot plane. This variance response for a given range of focal positions is shown in Fig. 4.

**Fig 4.**
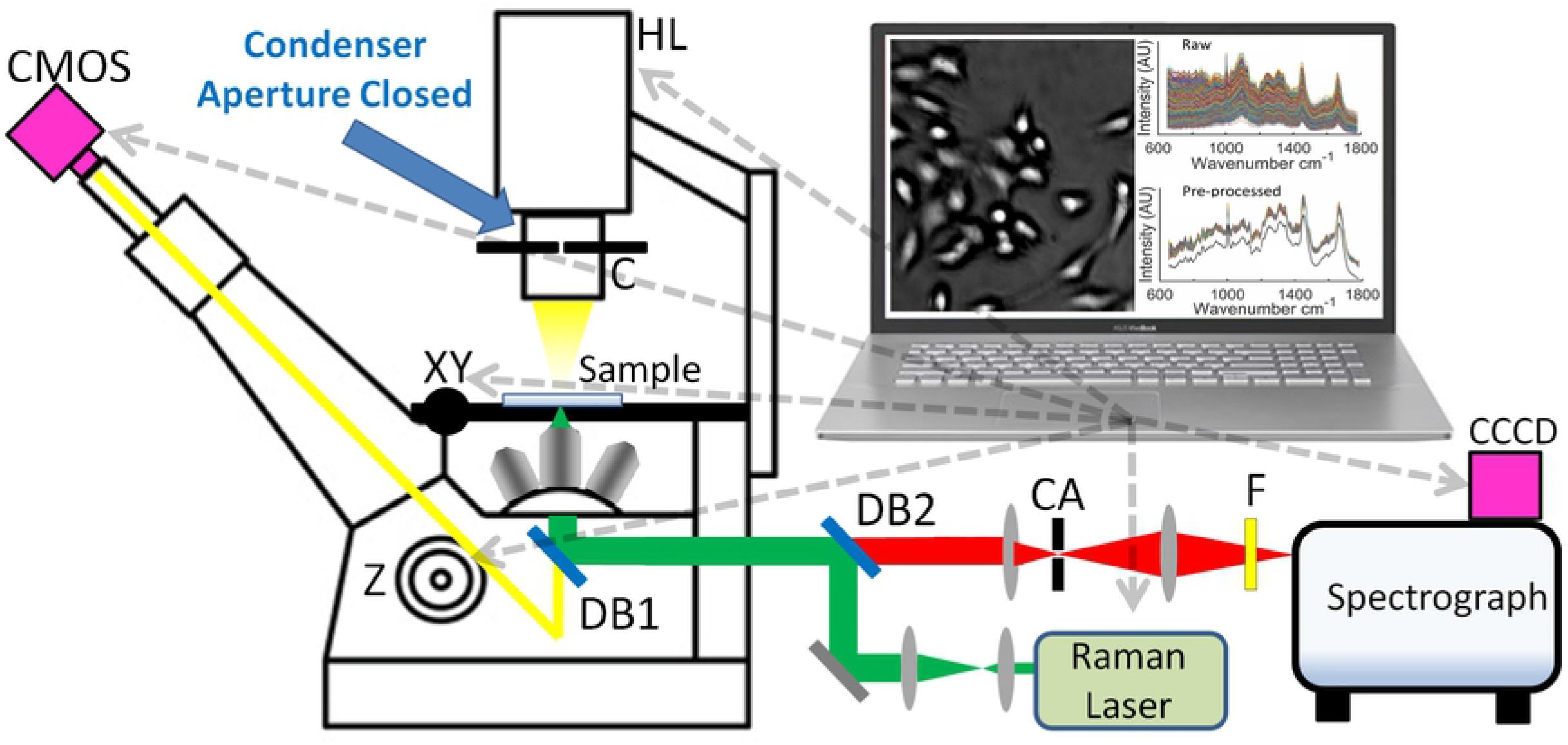
Variance profile shown on left side with points A and B corresponding to the sample positioned at the bright-spot plane and the focal plane of the microscope respectively - corresponding to images shown in Fig. 2 (c) and (a). On the right side a high-level flow chart is shown for the overall automated Raman cytology process, which includes moving the sample between these two planes.

The overall automation routine is illustrated in the flow-chart in Fig. 4 with a series of high-level steps. A more comprehensive low-level description is provided in the Supplementary Information, which corresponds directly with the source code provided in the online repository [26]. Once the bright-spot plane position is determined, it is relatively simple to acquire the positions of thousands of cells. A grid of overlapping images are recorded and stitched together using built-in ImageJ plugins. ImageJ analysis tools also allow for local maxima present within resultant image to be found and listed.

This list of coordinates includes thousands of cell nucleus positions, which can be further refined. Of these, a preset amount of the most isolated cells are sorted and filtered, removing the most clustered cells. By filtering for isolated cells, we improve the quality of the recorded spectra with respect to the targeting accuracy of the cell nucleus; as shown in the previous section, such clustered cells are more likely to be doubly targeted, particularly for the case of Thinprep slides. The refined list is then sorted using a nearest neighbour algorithm to limit stage movement, reducing the risk of drift and also speeding-up the overall acquisition process. Information from the image stitching process about image overlap relative to stage movement is used to create a coordinate transform, converting the list of cell positions from pixel coordinates to stage positions, which is described in more detail in the Supplementary Information. Once an offset for the laser spot position is included, and the stage is moved into the focus plane, the cells can be targeted by the Raman laser, and spectra can be recorded. The final step in the process is the removal of the baseline and glass spectrum component in these spectra as described in more detail in Section 3.5.

## 3 Materials and methods

### 3.1 Sample Preparation

#### 3.1.1 HeLa cell culture for fluorescence vs white-spot comparison

HeLa cells were cultured in DMEM media (Sigma-Aldrich) supplemented with 5% fetal bovine serum. Flasks were maintained in a humidified environment at 37°C and 5% CO_2_. When the cells reached 90% confluency, the culture media was removed, and the cells were washed with sterile PBS. 2mL of trypsin-EDTA solution (0.25%) was then added and the cells were incubated at 37°C for 5 minutes until the cells were detached. 8mL of DMEM media was added to the flask to neutralise the trypsin, and the entire contents were transferred to a sterile falcon tube. The cells were centrifuged for 5 minutes at 1500 rpm. The supernatant was removed, and the cell pellet was resuspended in fresh DMEM media. The cells were counted using a haemocytometer. After counting the cells, cells were prepared (1) for live cell imaging, and (2) using the ThinPrep standard; (1) For the live cell study, HeLa cells were seeded at 110^5^ cells/mL in a 12 well cell culture dish. The next day when the cell had reached 70% confluency, 2 drops of NucBlue™ Live ReadyProbes™ Hoechst 33342 (Thermo-Fisher) were added to each well. The cells were incubated for 20 minutes before imaging; (2) For ThinPrep, 500,000 cells were transferred to a new falcon tube and was centrifuged at 1500 rpm for 5 minutes. The cell pellet was resuspended in 20mL of PreservCyt (Hologic) and incubated for 20 minutes at room temperature. The cell suspension was loaded into the ThinPrep 2000 (Hologic) machine and the cells were transferred onto a glass slide. No coverslips were applied.

#### 3.1.2 Bladder Cancer cell lines for automated Raman cytology

High (HT1197; ATCC) and low (RT112, Sigma-Aldrich) grade urinary bladder carcinoma epithelium cells were cultured in 1:1 mixture of DMEM and Hams-F12 medium supplemented with 5% fetal bovine serum and 2 mM LGlutamine. Flasks were maintained in a humidified atmosphere with 5% CO_2_ at 37°C. When the cell lines reached 80% confluency, the culture medium was removed, and the cells were rinsed with sterile PBS. Trypsin-EDTA (0.5%) was added to the flask, which was incubated at 37°C until the cells had completely detached (not exceeding 15 min). An equal volume of 5% serum-containing medium was added to the flask to neutralise the trypsin enzyme. The entire contents of the flask was transferred into a sterile container, and centrifuged at 1200 rpm for 5 min. The supernatant was removed, and the cell pellet was resuspended in fresh medium. This solution was centrifuged at 1200 rpm for 5 min, the medium decanted, and resuspended in 1 ml PBS. This step was repeated and the cell pellets were resuspended into a vial containing 20 ml of a methanol based fixative (PreservCyt; Hologic, USA), and left at room temperature for 15 min. The vial was inserted into a ThinPrep 2000 (T2; Hologic, USA) machine, and the cells were transferred a CaF_2_ (Raman Grade; Crystran, UK) slide, or a glass slide (ThinPrep slide; Hologic, USA). No coverslips were applied.

### 3.2 Fluorescence Microscopy

Fluorescence images were recorded using an IX81 microscope attached to an MT20 illumination system with a 150 W Xenon arc burner and a filter set providing excitation at 358 nm and emission at 461 nm. Images were recorded on a low-noise CCD detector (QIClick™, QImaging, UK) and an Olympus LUCPlanFLN 40X/0.6NA microscope objective was used record the images, with variable coverslip correction 0-2 mm. Fluorescence images were captured with an acquisition time of 0.5 s and matching brightfield images were recorded with an acquisition time of 0.02 s. For the case of the live cells, the coverslip correction was set to 1 mm, while for ThinPrep it was set to 0 mm.

### 3.3 Automated Raman Optical System

Raman spectra from the cells described in Section 3.1 were recorded using a custom-built Raman micro-spectroscopy system, which is illustrated in Fig. 3. This system employs a 150 mW laser with a wavelength of 532 nm and a coherence length ≈100 m (Torus, Laser Quantum), which is driven by a power supply unit (mpc3000, Laser Quantum) that is controllable over an RS232 cable using Serial Commands using the Micro-Manager ‘freeserialport’ device adapter. The system also employs a spectrograph (Kaiser, Holospec f/1.8i) operating with a 25 μm slit and a holographic grating (HSG-532-LF). The spectrum is recorded using a low-noise cooled CCD camera (**CCCD**, DU920P-BEX2-DD, Andor) with 1024 x 256 pixels, of size 26 x 26 μm cooled to −80 °C and operating in full-vertical-bin mode. In this configuration, the camera provides a read noise standard deviation of 4 electrons/spectral sample, and a mean dark current of 0.0512 electrons/second/spectral sample. The camera was contolled using the Micro-Manager Andor device adapter. Together, the spectrograph and CCD provide a bandwidth of −34-2517 cm^−1^ and an average resolution of 5.48 cm^−1^.

Included in the spectrograph housing is a holographic notch filter (Kaiser; HSPF-532.0) providing an optical density of 6 at the laser wavelength and a spectral bandwidth of 350 cm^−1^. The laser and spectrograph were coupled into a fully automated inverted microscope (IX81, Olympus) controlled via the IX2-UCB control box, which allows for electronic control of the objective focus position (**Z**) and the white-light Halogen-Tungsten lamp (**HL**). This control box can be controlled using an RS-232 cable using the Micro-Manager Olympus IX81 device adapter. The microscope includes a closed-loop high precision stepper motor translation stage (**XY**, 96S108-O3-LE, Ludl) with a linear encoder, which provides repeatability of 0.25 μm and a resolution of 100 nm. The stage is driven by a control system (MAC 5000, Ludl), which be controlled using an RS-232 cable using the Micro-Manager Ludl device adapter. A 40x microscope objective, with numerical aperture of 0.75 (UMPlanFl, Olympus), is used to image the spectral irradiance to a 100 μm confocal aperture (**CA**, P100D, Thorlabs), which isolates the signal from the cell nucleus in three dimensions, and minimises background noise from the glass slides, as well as from optical elements in the system. The system is designed to provide a spatial resolution of ≈2-3 μm^2^ from the cell nucleus. A long pass filter (**F**, LP03-532RU-25, Semrock) and a dichroic beamsplitter (**DB1**, LPD-01-532RS, Semrock) are also used to filter the laser wavelength from reaching the spectrograph, while transmitting the longer Raman scattered wavelengths. A dichroic short pass filter (**DB1**, 69-202, Edmund Optic) permits imaging of the sample to a digital camera (**CMOS**, MU300, AmScope). All spectra were recorded using the Andor Solis software plugin for Micro-Manager. The system was wavenumber calibrated using a polymer standard as described in [33]. No intensity calibration was performed for this experiment since all spectra were recorded from the same system. The aperture in the microscope condenser (**C**, U-UCD8, Olympus) was closed to a minimum in order to maximise the spatial coherence of the illumination.

### 3.4 Spectral acquisition

The automation process described in Section 2 is capable of recording a large number of spectra. In total, spectra were recorded from 577 different HT1197 cell nuclei deposited on CaF_2_ (for the purpose of removing the glass spectrum as described in the following section), 6426 different HT1197 cell nuclei deposited on glass, and 7499 different RT112 cell nuclei deposited on glass. An acquisition time of 10 s was used to record a single spectrum from within the nucleus of each cell. In addition 20 spectra were recorded from the ThinPrep glass substrates, each of 30 s acquisition time, which is required during pre-processing as described below. Each recorded raw spectrum contained 1024 samples in the range −34-2517 cm^−1^, from which 451 samples were extracted for further inspection corresponding to the fingerprint region 600-1800 cm^−1^.

### 3.5 Pre-processing of Raman Spectra

Raw spectra were first input to a cosmic ray removal algorithm [34]; this algorithm is capable of removing cosmic rays from a spectrum by identifying a closely matching spectrum in the dataset, and replacing each cosmic ray with the corresponding wavenumber intensity values in that matching spectrum. It is, therefore, not necessary to apply the commonly used double acquisition cosmic ray removal method to obtain a closely matching spectrum [35], which was found to be problematic for the high-speed spectral acquisition applied in the automation process.

Cosmic ray removal was followed by an Extended Multiplicative Signal Correction (EMSC) algorithm [24, 36, 37], which removes the variable background signal from each spectrum. The EMSC algorithm estimates this background using an N-order polynomial (to remove the baseline signal that results from the cells auto-fluorescence) and also, if required, the background signal from the substrate. Briefly described, the EMSC algorithm applies a least squares fit to (i) a low-noise, contaminant-free reference Raman spectrum from a cell; (ii) an N-order polynomial; and (iii) a reference spectrum taken from the substrate; this is required for the case of glass substrates but not for the case of Raman-grade CaF_2_. The algorithm returns the weight of (i), which enables normalisation of the spectrum relative to the reference, as well as the total background made up of the appropriately weighted substrate spectrum plus the polynomial. The EMSC-corrected spectrum, *X*, is given by:

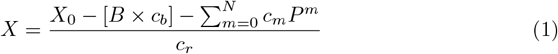

where *X*_0_ is the raw data, *B* is the reference spectrum of the substrate, *c_b_* is the weight of the reference substrate spectrum, *P^m^* denotes the *m^th^* order of the polynomial, *c_m_* is the corresponding polynomial coefficient, and *c_r_* is the weight of the cell reference spectrum, *R.* In summary, *X*_0_ can be described as the linear (weighted) superposition of *R, B,* and P. It has been shown that the use of a high-order polynomial does not result in over-fitting with the EMSC algorithm [37]. For this study, a fifth-order polynomial was used in the EMSC-correction algorithm for all datasets.

The reference cell spectrum provides the basis for all of the spectra to be fitted; the reference spectrum used here is the mean spectrum of the highest quality 50 spectra from the HT1197 dataset recorded on the CaF_2_ substrate. No smoothing was applied in this case. The background signal from Raman-grade CaF_2_ substrates are flat in the fingerprint region [38] and, therefore, this substrate was selected from which to obtain the high-quality reference cell spectrum to be used in the EMSC-correction of the spectra recorded on the glass substrates. In order to remove any potential bias, the same reference spectrum was used for the EMSC-algorithm applied to process the spectra of all cells deposited on glass. It has been demonstrated that equivalent results, in terms of the multivariate statistical analysis that follows, will be obtained when using significantly different reference spectra, so long as the same reference spectrum is applied in EMSC-correction of all datasets. [**?**] Also input to the EMSC-correction algorithm is a glass spectrum, which is the mean spectrum of the spectra recorded from the substrate followed by Savitsky-Golay smoothing using a polynomial order of 3 and a window size of 9.

Following EMSC correction, the spectra were denoised using a Savitsky-Golay based algorithm [39] using a polynomial order of 3 and a window size of 7. The resulting datasets of cell spectra were filtered in order to remove lower-quality spectra. This was achieved by removing all spectra that provided Pearson correlation coefficient of less than 0.99 with respect to reference cell spectrum, which resulted in a culling of approximately 50% of the data. A similar approach has previous been applied, albeit with a lower coefficient value, in order to extract spectra with high signal-to-noise ratios from a large dataset [15]. The results of the various pre-processing steps described in this section are presented in Section 4.

### 3.6 Multivariate Statistical Classification

In order to comprehensively evaluate the capability of the automated system to accurately classify the low- and high-grade urinary bladder carcinoma epithelium cells, and to elucidate the underpinning differences in their biochemical composition, a range of machine learning classification techniques were tested to discriminate the two spectral datasets, following the pre-processing steps outlined in Section 3.5. The algorithms considered were as follows: Linear Discriminant Analysis (LDA), Quadratic Discriminant Analysis (QDA), k-Nearest Neighbours (kNN), Random Forest (RF) [40], Support Vector Machine (SVM) [41] and Partial Least Squares (PLS) [42]. The classifiers were combined with two pre-processing steps, namely: Principal Component Analysis (PCA) [43] and Marginal Relevance (MR) for wavelength selection [44]. PCA obtains lower dimensional projections of the data in the feature space. The new features (Principal Components - PCs) represent directions in the observation space along which the data have the highest variability. In contrast, Marginal Relevance (MR) is a criterion that ranks each wavelength in order of their capability to discriminate between the classes. The MR score for each wavelength is the ratio of the between-class to within-class sum of squares. In this approach, each wavelength is considered independently of others and neighbouring wavelengths have similar MR scores.

All of the analysis was done in **R**, a free software environment for statistical computing and graphics [45]. LDA and QDA are implemented using the **MASS** [46] package and require no parameter tuning. PLS is implemented in the package **pls** [47] and the number of principal components was set to 15. SVM is implemented in the package **kernlab** [48]. Gaussian kernel was used with the bandwidth parameter value set at an empirical estimate suggested by [49]. kNN and RFs are implemented in the packages **class** [46] and **ranger** [50] respectively. The values for the tuning parameters of these models were obtained by cross-validation of the training set.

Ten-fold cross-validation was used to estimate the performance of the models on new data. In this application, PCA was carried out on the training set of each cross-validatory split of the data. Three PCs were used as input features to LDA, QDA and kNN classifiers. Marginal relevance (MR) criterion is implemented in the **R** package **BKPC** [51]. For all datasets, the highest scoring features from 7 regions were taken as inputs into the following classification algorithms: LDA, QDA, kNN, RF and SVM. RF, SVM and PLS were also trained on all wavelengths without any dimension reduction pre-processing steps.

## 4 Results

Following application of the Pearson correlation coefficient, the two spectral datasets of 6426 HT1197 (high-grade bladder cancer) cell spectra, and 7499 RT112 (low-grade bladder cancer) cell spectra, are reduced in number to 3583 and 3701 cell spectra, respectively. This corresponds to retention of 56% and 49%, respectively. The raw spectra corresponding to these two culled datasets are shown in Fig.5 (a) and (b), following cosmic rays removal. It is clear from the figure that there are significant differences across the dataset of raw spectra; this includes a variation in the minimum value of the different spectra resulting from changes in the camera temperature throughout the experiments, which affects the mean camera dark current from one recording to the next. Also noticable is a difference across the dataset in the relative amplitude of the various spectra, which results from slight variation in the focussing of the MO from cell to cell, as well differences in the morphology of the cell nuclei. In Fig.5 (c) and (d) the same two datasets are shown following pre-processing to remove the slightly varying baseline, and the glass spectrum as described in Section 2.2. Also shown in these figures are the mean spectra of the pre-processed datasets as well as the standard deviation of the pre-processed datasets around this mean value. It is clear from these results that the glass signal, most present in the raw spectra within the 1050–1150 cm^−1^ region has been reduced significantly.

**Fig 5.**
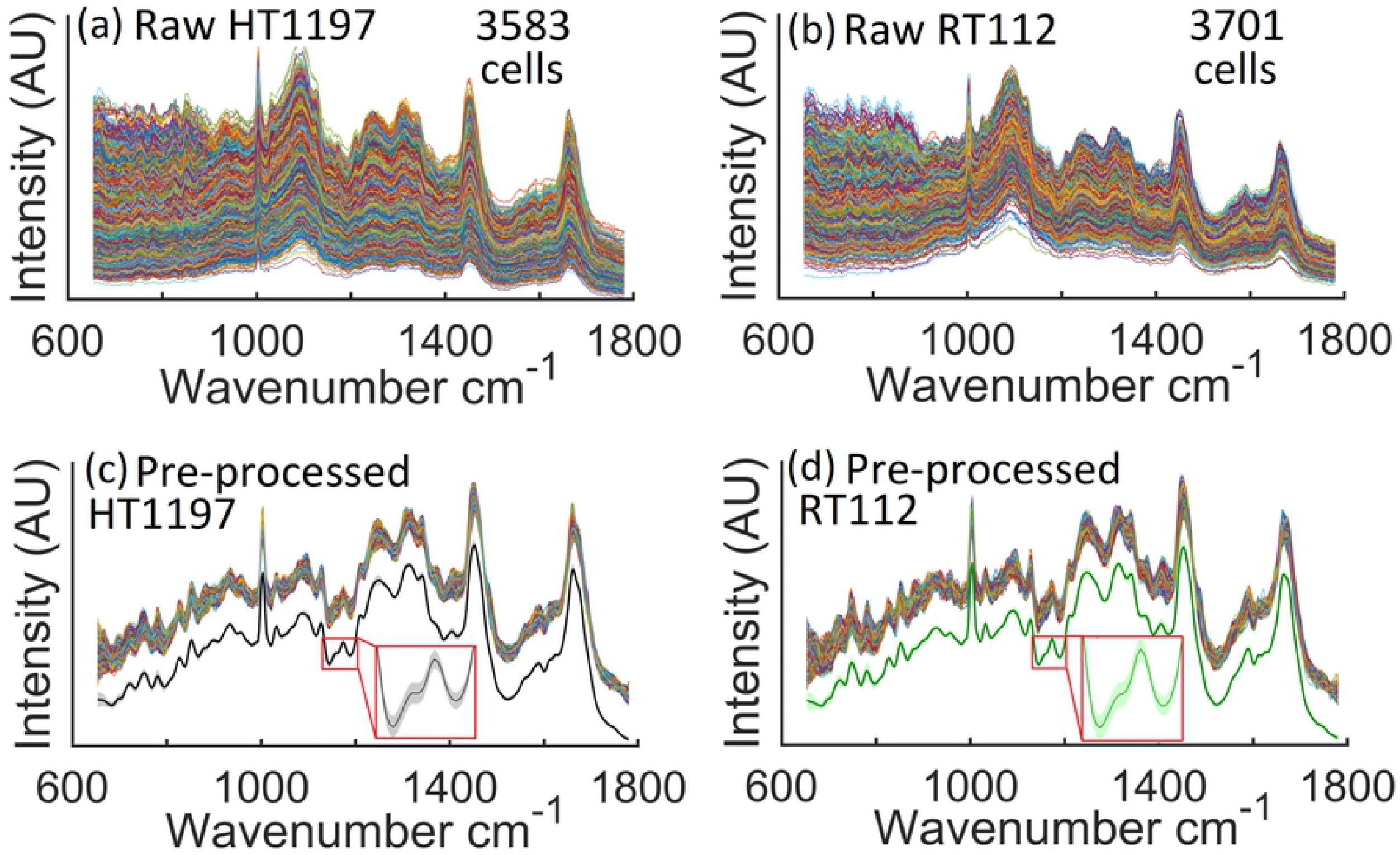
Raw and processed data from high- and low-grade bladder cancer cell lines. Raw spectra recorded using the proposed automated Raman micro-spectrometer, following cosmic ray removal; (a) 3583 spectra taken from individual HT1197 cell nuclei; and (b) 3701 spectra taken from individual RT112 cell nuclei; (c) The pre-processed HT1197 dataset, below which the mean spectrum is shown with a black line around which the standard deviation of the dataset is shown using a shaded grey colour; (d) The pre-processed RT112 dataset, below which the mean spectrum is shown with a green line around which the standard deviation of the dataset is shown using a shaded light green colour.

In Fig. 6 the results of Principal Components Analysis applied to the two pre-processed datasets. In Fig. 6 (a) a scatter plot is shown for the scores of the first two principal components and in Fig. 6 (b) the corresponding loadings are shown. It is clear that the two datasets separate well over these first two components. Inspection of these two principal components reveals several peaks that have previously been identified in the study of bladder cancer with Raman spectrocopy: 680 cm^−1^ (DNA), 789 cm^−1^ (DNA), 1003 cm^−1^ (Phenylalanine), 1170 cm-1 (Tyrosine), 1303 cm^−1^ (CH_3_, CH_2_, lipids, DNA), 1370 cm^−1^ (DNA, lipids), 1220 – 1300 cm^−1^ (Amide III, Collagen) [6,52] as well as peaks at 1093 cm^−1^ (PO2 stretching, DNA/RNA), 750 cm^−1^ (Tryptophan), 1310 cm^−1^ (Guanine), 1340 cm^−1^ (Adenine), and 1580 cm^−1^ (Tryptophan, DNA, phenylalanine), [5, 11, 52] and two new peaks that have not previously been identified in the analysis of bladder tissue: 1424 cm^−1^ (Deoxyribose) and 1490 cm^−1^ (DNA). [53]

**Fig 6.**
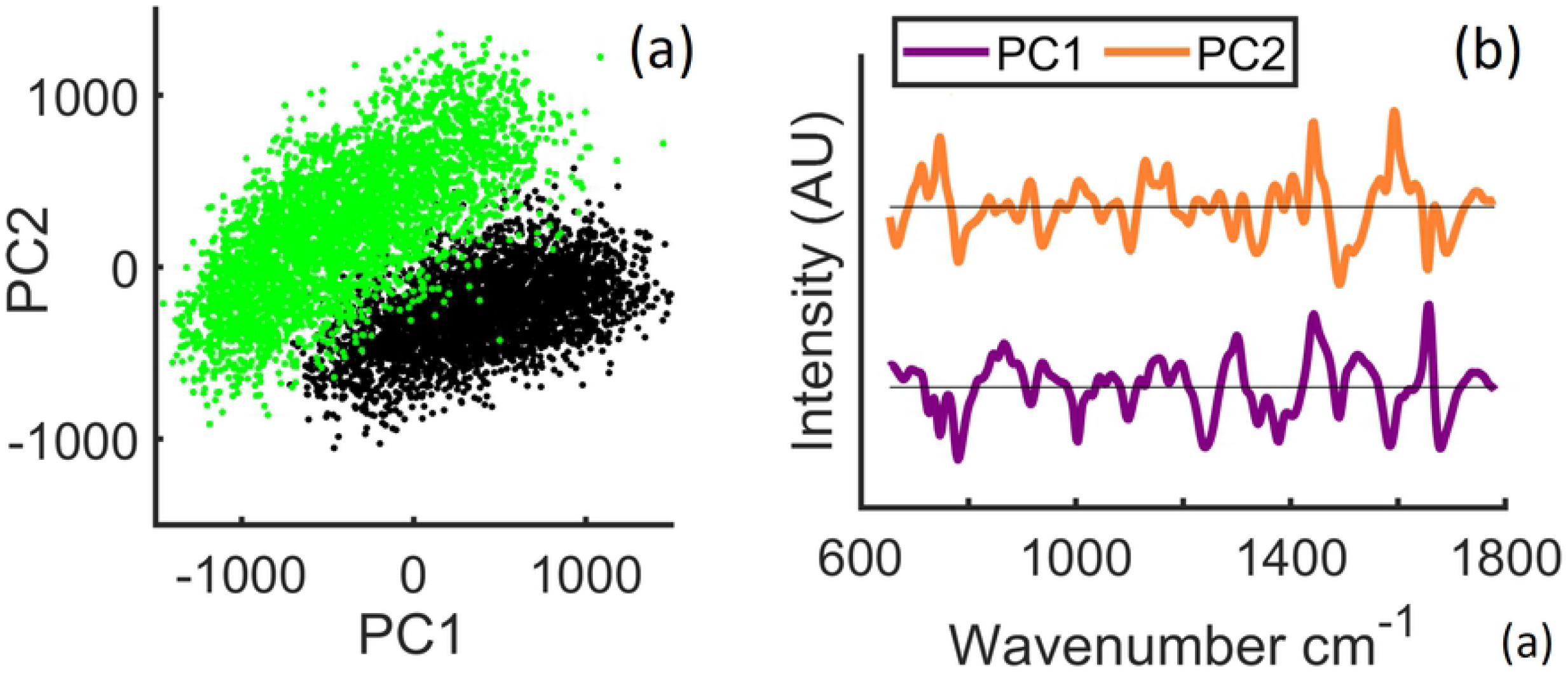
Multivariate statistical analysis (i) PCA scatter plot; (ii) First two principal components

Classification accuracy, sensitivity, and specificity for the eleven classification approaches are given in Table 1, and Fig. 7 shows box-plots of classification accuracy over the test sets in ten-fold cross-validation for the eleven classifiers. The comparative analysis suggests that classifiers PLS, SVM, and RF, with no (statistical) pre-processing steps, consistently perform better for classification of these datasets. MR seems to be more effective than PCA in selecting the features. We note that these sensitivies and specificities are the highest reported to date in the literature in the classification of high- and low-grade bladder cancer cell lines, likely owing to the increased reproducibility of the recording process as well as the increased dataset size.

**Table 1.** Classification accuracy, sensitivity, and specificity for the eleven classifiers: LDA, QDA and kNN applied after pre-processing with PCA and MR. RF and SVM were trained after pre-processing with MR and on the entire data without any pre-processing for dimension reduction. PLS was applied without any pre-processing.

|  | Accuracy | Sensitivity | Specificity |
| --- | --- | --- | --- |
| SVM | 0.996 | 0.996 | 0.996 |
| RF | 0.990 | 0.991 | 0.989 |
| PLS | 0.995 | 0.996 | 0.994 |
| PCA_LDA | 0.960 | 0.952 | 0.969 |
| PCA_QDA | 0.964 | 0.962 | 0.965 |
| PCA_kNN | 0.966 | 0.960 | 0.971 |
| MR_LDA | 0.981 | 0.980 | 0.982 |
| MR_QDA | 0.982 | 0.976 | 0.987 |
| MR_kNN | 0.980 | 0.975 | 0.984 |
| MR_RF | 0.982 | 0.980 | 0.984 |
| MR_SVM | 0.983 | 0.980 | 0.985 |

**Fig 7.**
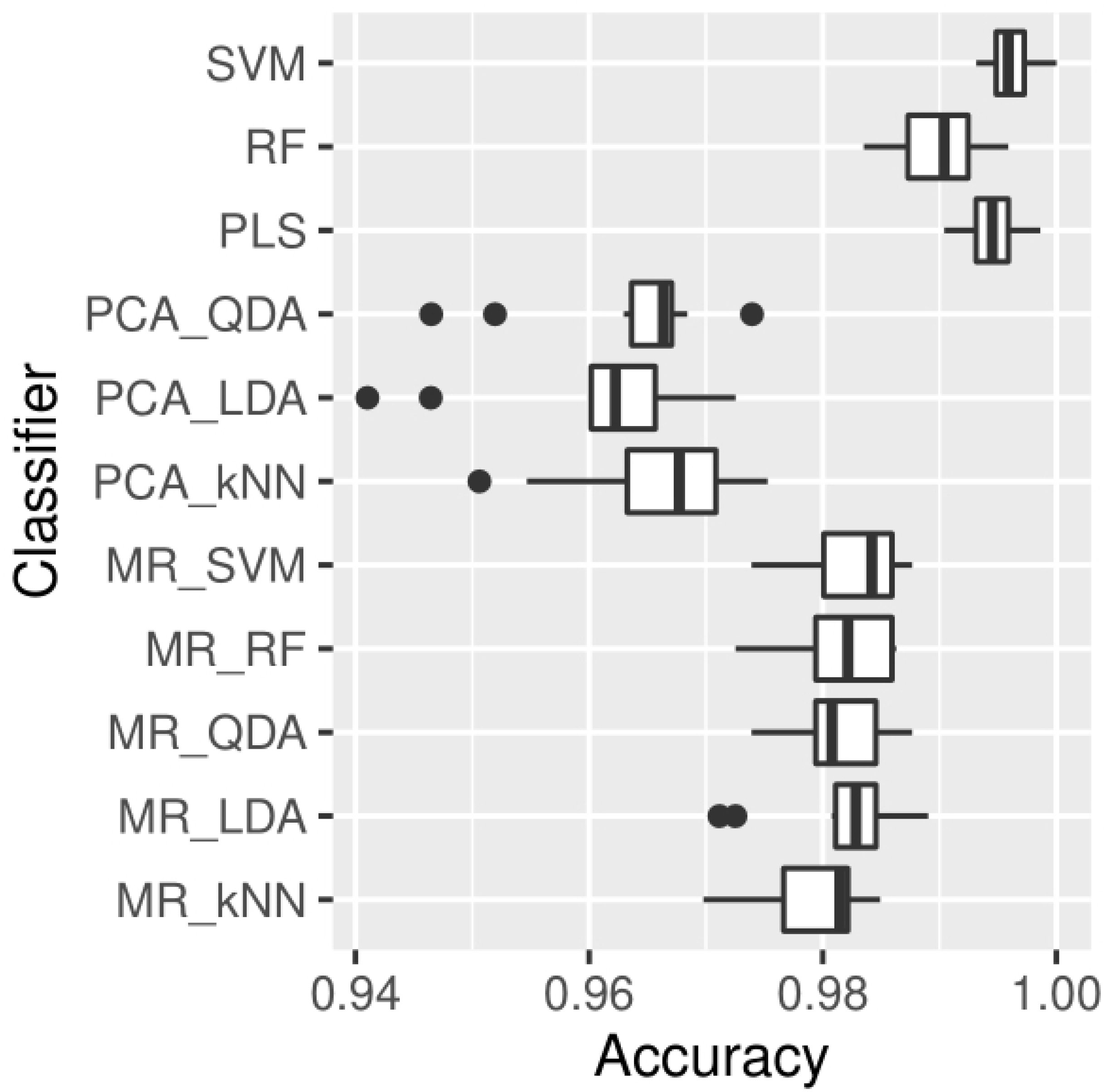
Classification accuracy of the ten test sets in ten-fold cross-validation for the eleven classifiers.

## 5 Discussion

The automated Raman cytology system presented here is is focused on recording spectra from the unlabelled nucleus of epithelial cells. Automated targeting of the nucleus is a key differentiator of the proposed technology with other automated Raman cytology systems, which have recently been proposed and which target the centre of the cell mass [15, 16]. The basis of our approach is to iterate the focus of microscope between two planes, the traditional image plane at which Raman spectra are recorded, and the ‘bright-spot plane’ some tens of micrometers from the image plane, where the ‘micro-lens’ effect of the cell nuclei focuses the partially coherent microscope illumination and produces bright spots approximately co-located with the cell nuclei, thereby facilitating subsequent RS. Our approach builds on the work of Drey et. al. [29], who first identified the phenomenon and utilised it for the purpose of cell counting of live cell cultures. The ‘focal-length’ of the nucleus, i.e. the distance between the microscope focal plane and the ‘bright-spot plane’ appears to depend on the spatial variation in cell-thickness with a thicker, rounder morphology resulting in the focusing of light several micrometers behind the sample plane; and a thinner, flatter morphology focusing the light several tens of micrometers behind the sample plane.

We have shown that in addition to live adherent epithelial cells, the method can also be applied to identify the nucleus of cells prepared using the ThinPrep standard, which includes the use of specific fixatives and glass slides. Appending such a widely established clinical practice for preparing uniform patient samples for subsequent pathological classification, is an attractive means for advancing Raman cytology into the clinic as an assistive tool for cyto-histopathologists. HeLa cells prepared using ThinPrep, and compared with the same cells growing in medium, revealed a stark difference in morphology with the former appearing significantly thicker and having a smaller footprint. Interestingly, this translated to a significantly shorter nucleus ‘focal-length’. Nevertheless, identifying the nucleus could be achieved for either method of cell preparation, which indicates that the proposed automated Raman cytology system could be applied both to growing cell lines for the purpose of basic research, as well as to patient cytology samples prepared using the ThinPrep standard. Over both methods of cell preparation we have found an accuracy of nucleus identification for approximately 75% of cells. We believe this could be improved upon by using a colour filter in the illumination lamp to increase bright-spot contrast as in [29], or for the case of live cells, to immerse the cells in phosphate buffered solution before imaging, which was shown in [29] have the effect of swelling the cell body and significantly enhancing bright-spot contrast.

The overall throughput of the method is demonstrated at approximately 0.1 cell/sec, which is slower than the method proposed by Schie et. al. [15], which can record at a rate of approximately 1.4 cell/sec. However, this comparison must be qualified by the cell type that have been investigated by the two systems. In [15] the authors applied their system to lymphocytes, neutrophils, and monocytes, which are considerably thicker than the epithelial cells investigated in this paper, and which, therefore, produce a more intense Raman scattering, necessitating a shorter acquisition time. Furthermore, they use a significantly more powerful laser source of 400 mW, albeit at a longer wavelength of 785nm that will produce less Raman scattering [38]. Noticeably, they also use a Pearson correlation coefficient of 0.95 to cull their noisy spectra while we use of a coefficient value of 0.99. We have found that an acquisition time of 5 s (increasing the throughput of 0.2 cells/sec) could be used if we apply a value of 0.95. We believe that it may be possible to further reduce acquisition time by increasing the laser power, distributed over a larger area of the cell nucleus, and using advanced denoising methods based on machine learning. With these approaches, we believe it may be possible to achieve 1 cell/sec throughput, and this will be a subject of our future work.

Although we have demonstrated the applicability of the automated identification of the nucleus for both live cells and cells prepared using ThinPrep, we focus our experiments in Section 4 on bladder cancer cell lines using the latter. The application of the proposed system to clinical cytology samples is the primary motivation of our work. An important feature of our automated platform is the use of 532 nm excitation and the removal of the glass spectrum, which is unavoidable for clinical deployment of the technology. We have identified at least three clinical areas that could potentially benefit from the proposed technology, all of which have been shown to be diagnostically improved by Raman spectroscopy: The most common branch of cytology is the ‘Pap smear,’ used to screen precancerous cervical lesions; application of Raman to cervical cytology samples has received significant attention in the literature [7, 11–13]. Cytological inspection is also common for bladder cancer, whereby epithelial cells are extracted from urine, though this is often used as an adjunct to cystoscopy due to the low sensitivity (20%) for low grade carcinoma accounting for the majority of cases. RMS has been demonstrated to improve the sensitivity of urine cytology to ¿90% [3,4,6] and our group has actively researched a methodology for RMS to be integrated into the pathology lab [5]. Oral cancer is one of the most common cancers worldwide with tumours located around the tongue and mouth. Late stage oral cancer is straightforward to diagnose with histological analysis of tissue biopsy, but results in poor outcome. RMS has been demonstrated to successfully identify precancerous tissue by investigating epithelial cells [8,14]. All three of these clincial areas require only non-invasive procedures to retrieve cells, and although each of them benefit from improved classification using RMS, clinical adoption has been slow due to the slow throughput of Raman and issues with reproducibility. We believe that the system proposed here can solve these issues and facilitate clinical adoption.

A final point of note in this discussion is the development of the proposed automation routine using the Micro-Manager platform. This software is open source and readily applicable to any existing electronically controllable microscope and RMS systems that are commercially available. The programming scripts that we have developed are provided in an online repository [26] and can be downloaded and adapted with relatively little effort. We hope this approach will help to advance automated Raman cytology for clinical applications.

## 6 Conclusion

In this paper, we have demonstrated an automated Raman cytology system that is capable of targeting epithelial cell nuclei, for the primary purpose of improving the throughput and reproducibility of the clinical application of Raman cytology. Importantly, the system can be applied to both living adherent epithelial cells in medium for basic research, as well as cells prepared using the ThinPrep standard, an established cell fixation and deposition protocol for cervical, urine, and oral cytology, which have all previously been shown to be enhanced by Raman spectrocopy. Using two clinically relevant cell lines, a high- and low-grade bladder cancer cell line we demonstrate a cell throughput of 0.1 cell/sec and discuss approaches for further increasing this. The automation routine is developed using the open-source Micro-Manager software system and can readily be downloaded and adapted for existing RMS systems. We hope that this work will provide the much need throughput and reproduciblity to finally advance Raman cytology into routine clinical practice.

## Acknowledgments

This work is supported by Science Foundation Ireland under Grant Number 15/CDA/3667.

---

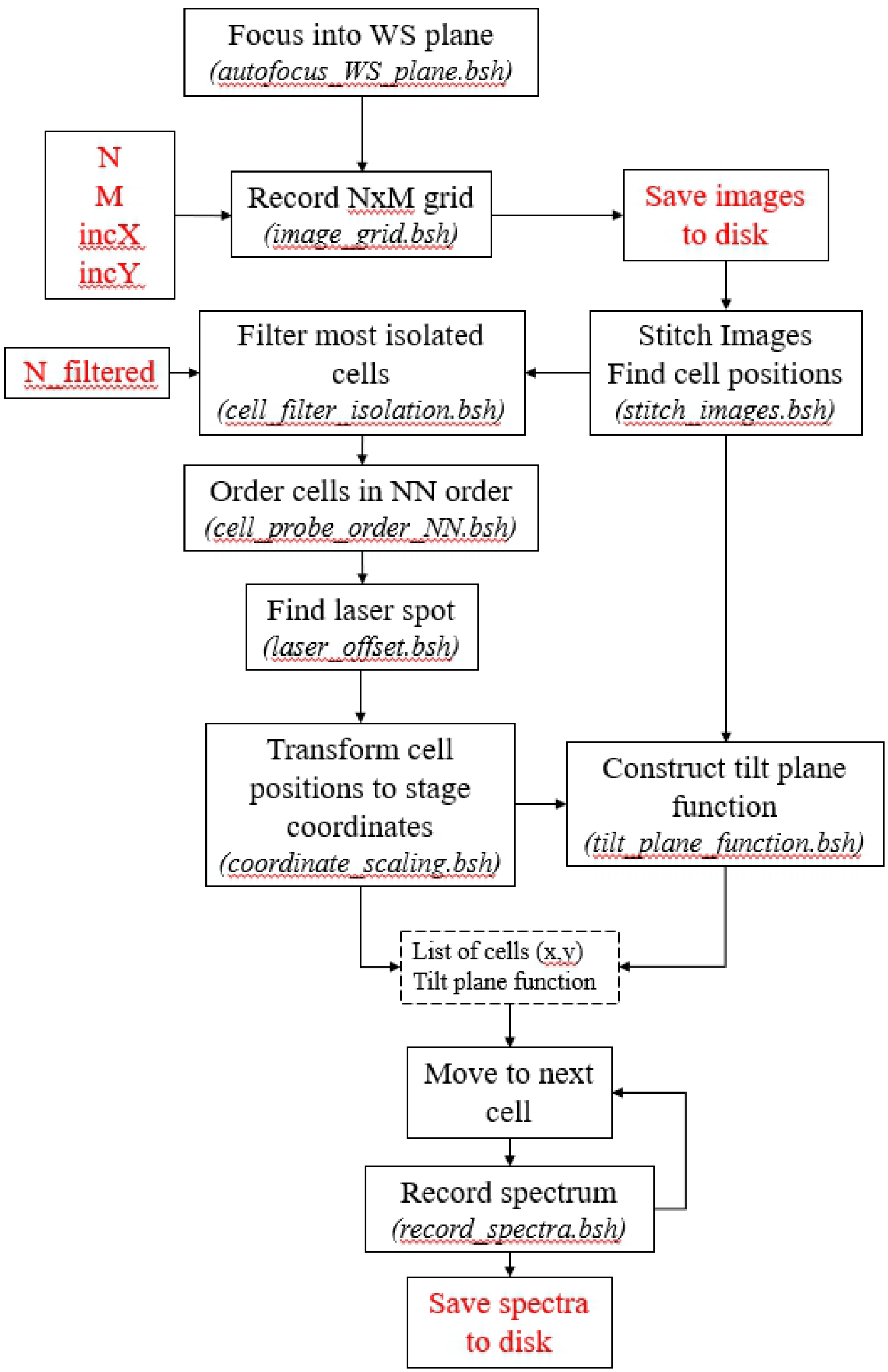

